# Overlooked neuroanatomical markers of face processing and developmental prosopagnosia in posteromedial cortex

**DOI:** 10.1101/2025.09.18.677204

**Authors:** Joseph P. Kelly, Ethan H. Willbrand, Xiayu Chen, Samira A. Maboudian, Benjamin J. Parker, Guo Jiahui, Lúcia Garrido, Zonglei Zhen, Bradley Duchaine, Kevin S. Weiner

**Author notes:** Corresponding author and lead contact: Kevin S. Weiner.

## Abstract

Recent functional imaging studies implicate human posteromedial cortex (PMC)—composed of the posterior cingulate cortex, the precuneus, and retrosplenial cortex—in face processing. Separately, anatomical studies have identified previously overlooked cortical folds (sulci) in PMC associated with higher-level cognitive abilities. Here, we tested whether these newly identified sulci support face processing in neurotypical individuals and individuals with developmental prosopagnosia (DP). After manually labeling 1,642 sulci in 164 hemispheres, we first identified a gradient of face selectivity along the anterior–posterior axis of PMC that was consistent across three samples, including DP individuals. Second, we discovered a new anatomical locus in PMC that differed structurally and functionally between neurotypical and DP individuals. Finally, data-driven analysis revealed that right-hemisphere PMC sulcal morphology was associated with face recognition ability. These findings reveal a sulcal network in PMC that supports face processing, and they identify the first structural neuroanatomical marker of face processing deficits in PMC.

## Introduction

The posteromedial cortex was once dubbed the “medial mystery parietal area.”^1^ Understanding its structure and function is of great interest to neuroscience. Functionally, the PMC is a site of peak metabolic activity within the cerebral cortex and serves as a connectivity hub for several large-scale brain networks.^2,3^ It supports numerous cognitive functions including memory, navigation, emotion, and executive control.^2,3^ Adding to this variety of PMC functions, recent studies have identified a “medial face area” in posterior PMC that is active during the recall and recognition of familiar faces.^4–8^ These studies are the most recent additions to a long-standing literature identifying face-selective areas throughout the human cerebral cortex, including parts of PMC,^9–33^ and they emphasize a specific role for PMC in the mnemonic processing of familiar faces. But these findings also raise several questions: How does the anatomy of PMC support face processing? Does the structure or function of PMC differ in populations with face processing deficits? Beyond its role in familiarity and memory processing, could PMC play a more fundamental role in the visual processing of faces?

Structurally, PMC is traditionally divided into the posterior cingulate cortex (PCC), the precuneus (PrC), and the retrosplenial cortex, but recent work has enriched this picture by identifying anatomical subdivisions within PMC that are consistent across species—including a “visual posterior precuneal region”—with distinct structural and functional connectivity profiles.^34,35^ Adding to these advances, recent studies have identified new cortical indentations (sulci) in PMC, a subset of which are (1) hominoid-specific, (2) associated with individual differences in memory and executive functioning abilities, and (3) especially vulnerable to age- and Alzheimer’s-related atrophy.^36–38^

Several studies have linked face processing to precise anatomical features in “core” visual areas of the face processing network such as the fusiform face area (FFA).^39–43^ But it is presently unknown whether anatomical features in “extended” areas of the face processing network, such as PMC, are associated with face processing—e.g., whether recently identified PMC sulci respond selectively to face stimuli, or whether their structural characteristics are associated with face recognition ability. It is also unknown if these sulci differ in clinical populations with face processing deficits, such as individuals with developmental prosopagnosia (DP; ∼2% of the population who have lifelong difficulty recognizing faces despite no history of brain damage).^44,45^ Establishing such links between individual anatomy and face processing ability may uncover precise areas of abnormal cortical development in individuals with DP. More broadly, it will advance our understanding of the neuroanatomical basis of face processing in the extended face processing network.

In the present study, we tested whether structural and functional features of PMC sulci are associated with face processing in neurotypical (NT) individuals and individuals with DP. To do so, we leveraged previously published datasets^40,46^ that include anatomical MRI, functional MRI, and behavioral data from NT (N=43) and DP (N=39) participants. To test the reproducibility of any identified structural-functional relationships, we independently analyzed data from the Human Connectome Project (HCP), leveraging the fact that (1) PMC sulci were identified in a subset of HCP individuals (N=71) by Willbrand and colleagues^36–37^, and (2) these HCP individuals participated in an experiment to localize face-selective regions.^47–49^

After manually labeling 1,642 PMC sulci in 164 hemispheres (Figure 1), we implemented a multimodal approach to test if PMC sulci were associated with face processing. We report three main findings. First, BOLD responses in recently defined^36–37^ sulci showed face selectivity in posterior PMC, but not anterior PMC. This result was consistent across NT and DP individuals and replicated in an independent dataset from the HCP. Second, we identified structural– functional differences between DP and NT individuals; a newly identified sulcus in ventral PCC was present less frequently and showed lower face selectivity in DP compared to NT individuals in the right, but not left, hemisphere. Third, a data-driven supervised learning approach revealed that three recently defined PMC sulci in the right hemisphere are associated with behavioral performance on a canonical face processing task. Together, these findings identify precise neuroanatomical features in PMC that support face processing, and they advance a growing literature identifying sulcal-functional couplings in face-selective regions across the cerebral cortex. They also provide the first evidence of anatomical differences in PMC between DP and NT individuals, establishing new targets for clinical research. Finally, they advance theoretical accounts of face processing by re-situating PMC within both the core and extended aspects of the human face processing network.^9,10^

**Figure 1:**
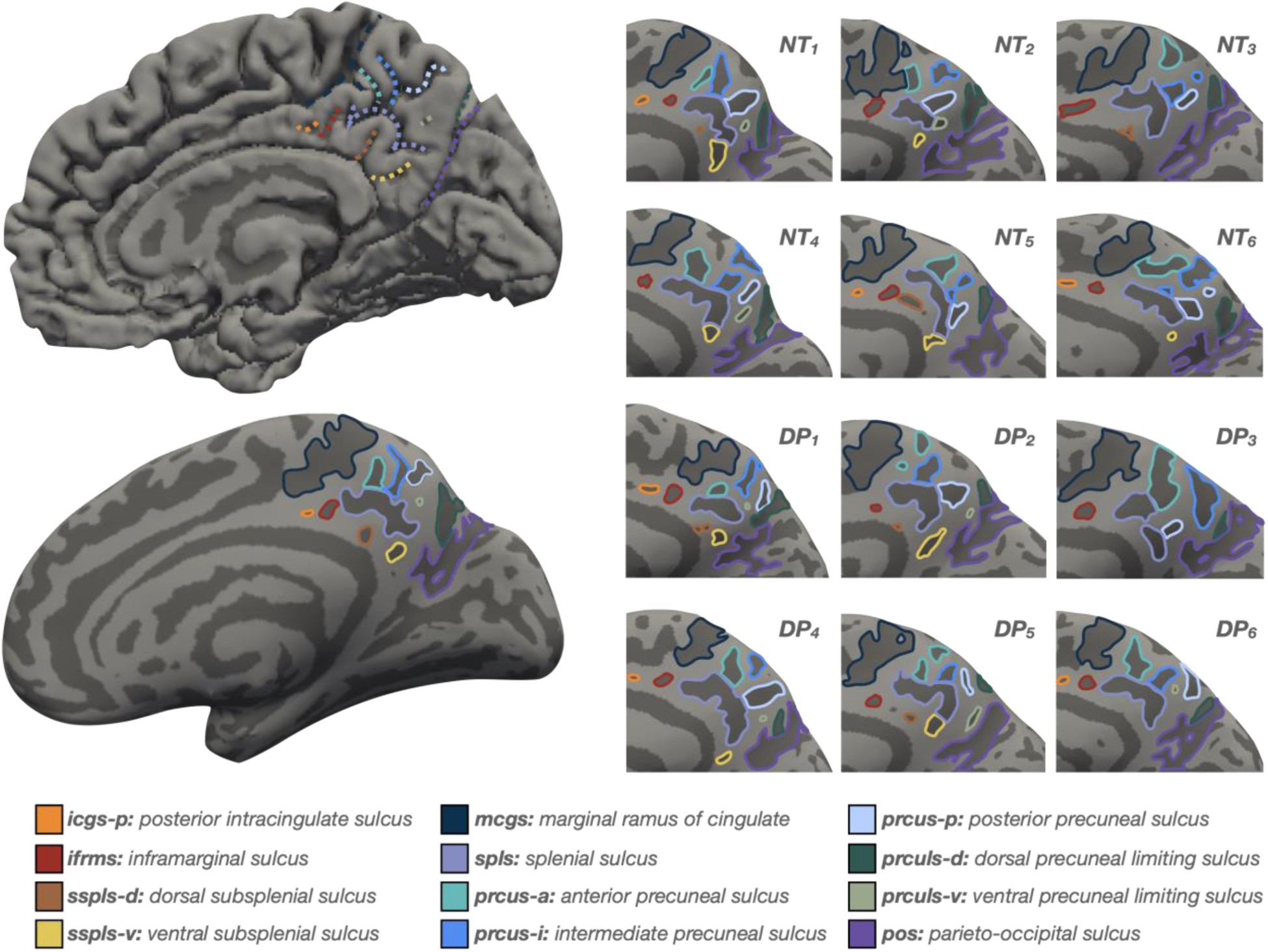
Manually defined PMC sulci in neurotypical controls (NT) and individuals with developmental prosopagnosia (DP). Top left: A cortical surface reconstruction of an individual right hemisphere (medial view). Sulci: dark gray; Gyri: light gray. Individual sulci are depicted by colored outlines (legend). Bottom left: The same cortical surface but inflated. Right: Twelve example hemispheres, six from NTs (top) and six from DPs (bottom). Left-hemisphere images are mirrored so that all images have the same orientation. Recently identified sulci include the icgs-p, ifrms, sspls-d, sspls-v, prcus-a, prcus-i, prcus-p, and prculs-v.^36–37^

## Results

### A sulcal-functional gradient of face selectivity along the anterior–posterior axis of PMC

To determine if structural and functional features of PMC sulci were associated with face processing, we first tested all PMC sulci for face selectivity in a subset of NTs (N=25) and DPs (N=22) who participated in a functional localizer fMRI experiment.^46^ We conducted a mixed-model ANOVA examining mean face selectivity [faces - objects; % signal change (as in Jiahui and colleagues)^46^] across factors of sulcus (all PMC sulci included), hemisphere (left, right), and group (NT, DP). This analysis revealed a three-way interaction between sulcus, hemisphere, and group [*F*(11,414) = 2.80; *p* = .002; η^2^*_G_* = .017]. Post hoc one-sample t-tests revealed two sulci in the posterior PrC (Figure 2a) that showed consistent face selectivity across both hemispheres and groups (prcus-i, prcus-p: *p*s ≤ .008; all one-sample tests Bonferroni-corrected for multiple comparisons). Further, post hoc pairwise contrasts revealed a sulcal–functional gradient of face selectivity across the anterior–posterior axis of the PrC that was consistent across hemispheres and groups: the prcus-p showed significantly greater face selectivity than the prcus-a across hemispheres and groups (*p*s ≤ .014; all pairwise tests Tukey-corrected for multiple comparisons), and the prcus-i showed greater selectivity than the prcus-a in both hemispheres of NTs (*p*s ≤ .019) (Figure 2b, left; see Supplementary Materials for results for all sulci).

**Figure 2:**
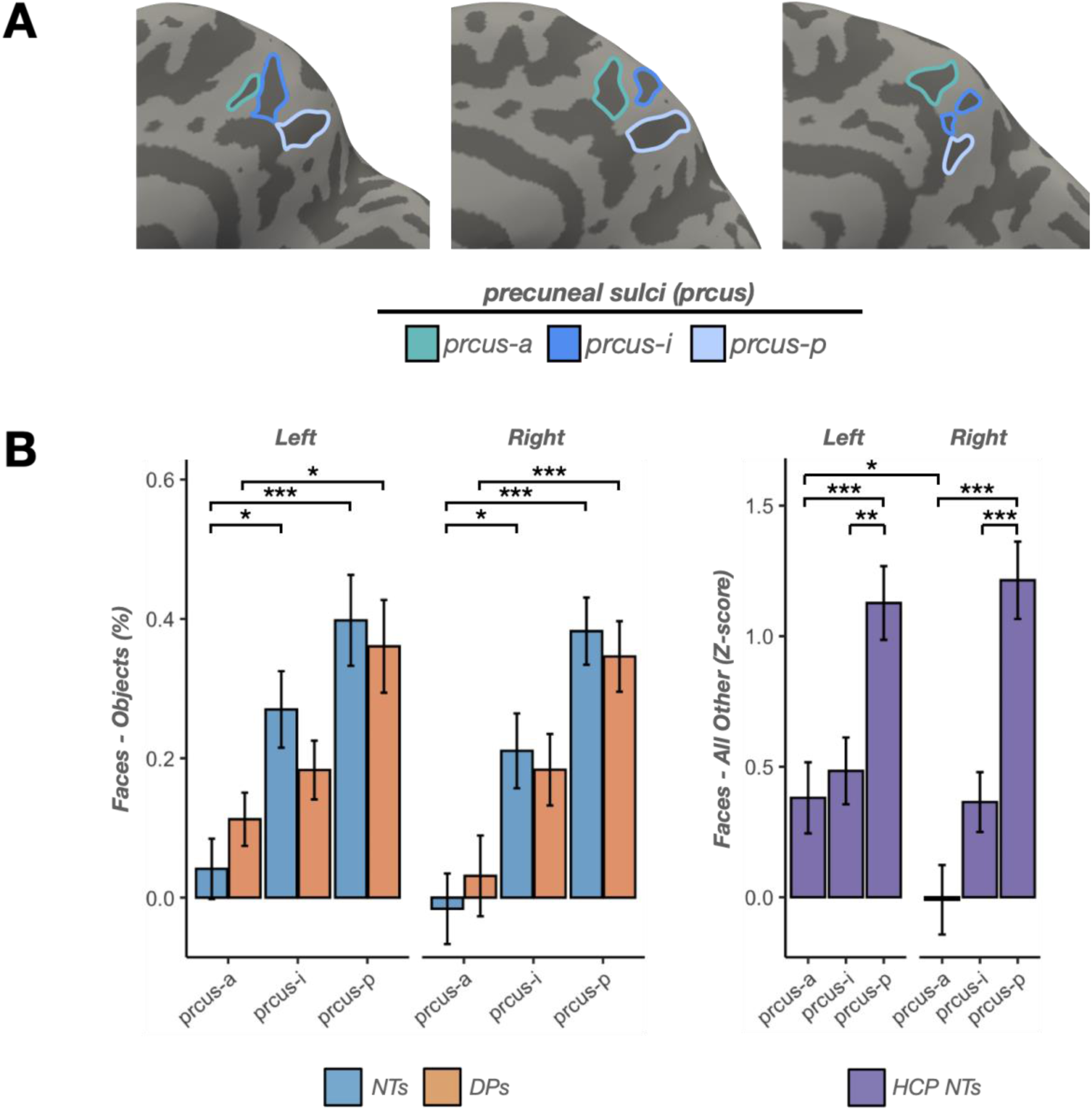
A sulcal-functional gradient of face selectivity along the anterior–posterior axis of PMC. (A) Inflated cortical surface reconstructions from example participants. Sulci: dark gray; Gyri: light gray. Individual precuneal sulci are outlined according to the legend. (B) Left: Face selectivity [faces - objects; % signal change (as in Jiahui and colleagues)^46^] of each precuneal sulcus (prcus-a: anterior precuneal sulcus; prcus-i: intermediate precuneal sulcus; prcus-p: posterior precuneal sulcus) in NTs (blue; N=25) and DPs (orange; N=22). Face selectivity increased along the anterior–posterior axis of the precuneus in both hemispheres and in both NTs and DPs, with no significant differences between groups. Right: Face selectivity [faces - all other categories; z-score (as in Chen and colleagues)^76^] in an independent dataset of NT participants from the Human Connectome Project (HCP, purple; N=71). The same increase in face selectivity was observed along the anterior–posterior axis of the precuneus in both hemispheres. Error bars in all plots indicate ±1 standard error. Significance values: *p<.05; **p<.01; ***p<.001.

To assess whether the same sulcal–functional gradient of precuneal face selectivity was present in an independent dataset, we ran the same analysis in a sample of 71 HCP NT participants (Dataset 3, STAR Methods) whose PMC sulci had been manually defined previously.^36–37^ A mixed-model ANOVA for mean face selectivity [faces - all other categories; z-score (as in Chen and colleagues)^76^] with the same fixed effects as above yielded an interaction effect between sulcus and hemisphere [*F*(11,649) = 4.46; *p* < .0001; η^2^*_G_* = .021]. Post hoc analyses confirmed findings from the first analysis, showing that the prcus-p was strongly selective for faces across both hemispheres in HCP NTs (*p*s ≤ .0001). Further, pairwise contrasts showed that the prcus-p had significantly greater face selectivity than the prcus-a and prcus-i in both hemispheres (*p*s ≤ .009; Figure 2b, right; see Supplementary Materials for results from all sulci). This result shows that the anterior–posterior gradient of PrC face selectivity observed in the NT and DP samples generalized to an independent sample of NTs. Critically, the stimuli and task used for this fMRI localizer contrast were completely different between the two datasets, indicating that the result is not stimulus- or task-specific. Taken together, these results identify a sulcal–functional gradient of face selectivity along the anterior–posterior axis of the PrC that (1) is consistent across hemispheres, (2) is present for both NTs and DPs, and (3) generalizes to an independent sample of NTs.

### The ventral sub-splenial sulcus (sspls-v) shows reduced incidence and face selectivity in developmental prosopagnosia

This is the first time that PMC sulci, including recently identified shallow sulci,^36–37^ have been characterized in individuals with DP. Some of these recently identified shallow sulci are present in every individual (e.g., ifrms), while others (e.g., sspls-v) are highly variable (Figure 3a; see Table S1 for incidence rates of variably present sulci). The presence or absence of small and shallow sulci has been linked to individual differences in cognitive abilities in both neurotypical individuals and individuals with neurodevelopmental disorders.^77–81^ The sspls-v is of particular interest in a study of developmental prosopagnosia because it is anatomically located in the same ventral-PCC region as the “medial face area” identified in fMRI and intracranial EEG studies; this region is active during the processing of familiar faces.^4–8,10^ We therefore used chi-squared tests to examine whether sspls-v incidence (% of hemispheres in which the sulcus is present) differed by group (NT, N=43; DP, N=39) or hemisphere (left, right). In the right hemisphere, DPs showed significantly lower sspls-v incidence than neurotypical controls (DP: 41.0%, NT: 67.4%; 𝝌*^2^*(1) = 4.75, *p* = .029), while left-hemisphere incidence was comparable between groups (DP: 76.9%, NT: 83.7%; 𝝌*^2^*(1) = 0.25, *p* = .619). Within DPs, the sspls-v appeared significantly less frequently in the right than left hemisphere (right: 41.0%, left: 76.9%; 𝝌*^2^*(1) = 8.96, *p* = .003), whereas neurotypical controls showed no significant hemispheric asymmetry (right: 67.4%, left: 83.7%; 𝝌*^2^*(1) = 2.27, *p* = .132).

**Figure 3:**
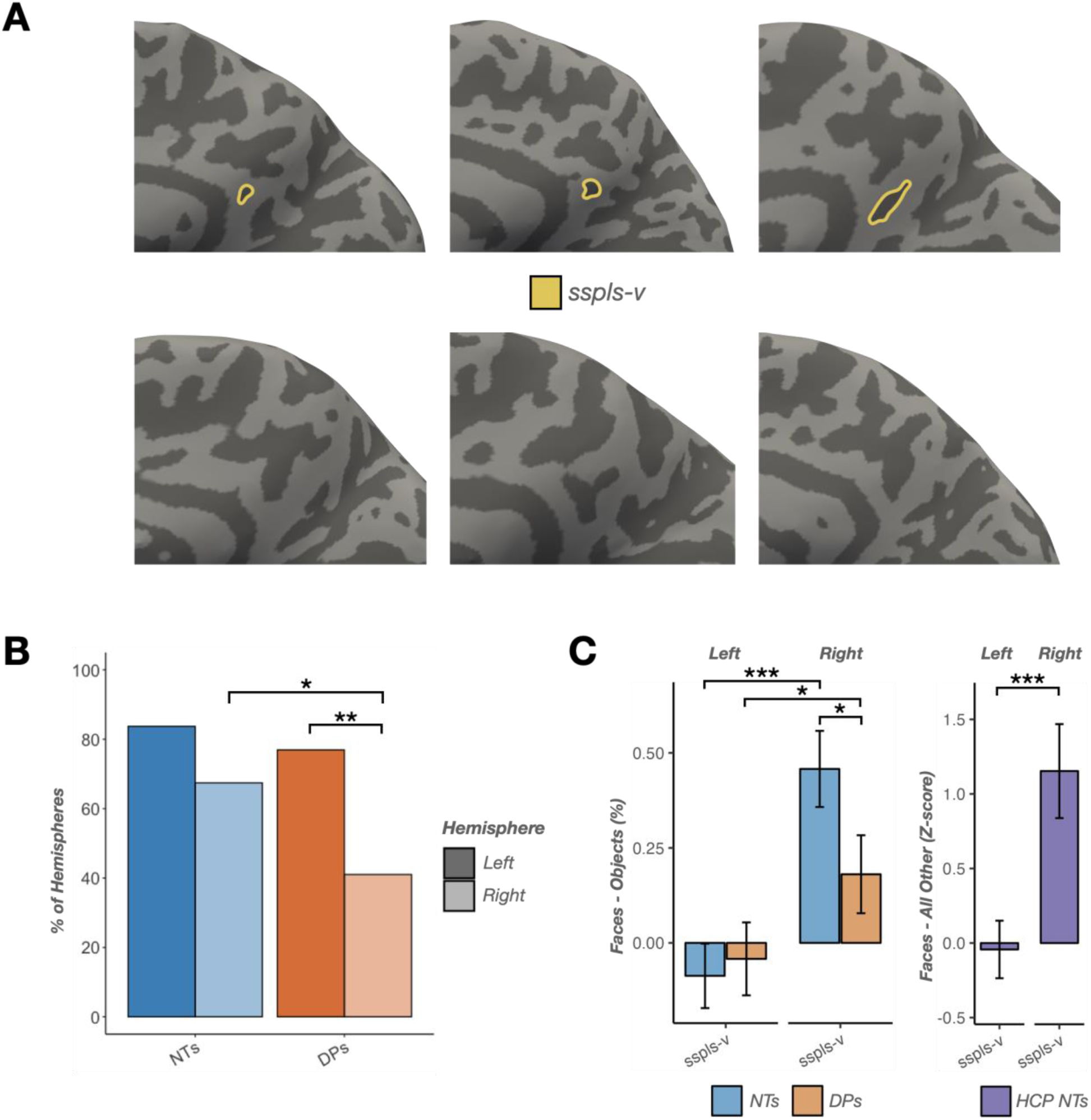
The ventral sub-splenial sulcus (sspls-v) shows reduced incidence and face selectivity in developmental prosopagnosia. (A) Top: Inflated cortical surface reconstructions from example participants with a ventral sub-splenial sulcus (sspls-v). Sulci: dark gray; Gyri: light gray. The sspls-v is outlined in yellow. Bottom: Three example hemispheres without an sspls-v. (B) Left: Sulcal incidence (% of hemispheres in which a sulcus is present) of the sspls-v in the left (dark) and right (light) hemispheres of NTs (blue; N=43) and DPs (orange; N=39). DPs showed significantly lower right-hemisphere incidence than NTs, and significantly lower right-hemisphere incidence than left-hemisphere incidence. (C) Left: Face selectivity (faces - objects; % signal change) of the sspls-v in NTs (blue; N=25) and DPs (orange; N=22). Face selectivity of the right-hemisphere sspls-v was significantly lower in DPs compared to NTs. Additionally, the sspls-v showed significantly greater face selectivity in the right hemisphere compared to the left in both NTs and DPs. Right: The right>left hemispheric effect generalized to an independent dataset of NTs from the HCP (purple; N=71). Error bars indicate ±1 standard error. Significance values: *p<.05; **p<.01; ***p<.001.

Face selectivity analysis revealed functional differences that parallelled the anatomical findings. Using the same mixed-model ANOVA run previously (NT, N=25; DP, N=22), we found that the sspls-v was face-selective in the right hemisphere of NTs (*p* ≤ .0001) but not in the left hemisphere (*p* = .289) nor in either hemisphere of DPs (*p*s ≥ .108). Between-group comparisons showed stronger face selectivity in NTs compared to DPs in the right hemisphere (*p* = .002) but not the left (*p* = .747). Additionally, both groups showed right>left face selectivity (NTs: *p* < .0001, DPs: *p* = .031), a finding that generalized to an independent dataset of 71 NTs from the HCP (*p* < .0001). Together, these converging anatomical and functional findings identify the sspls-v as a neuroanatomical site differentiating individuals with DP from neurotypical controls in both sulcal incidence and face selectivity (Figure 3).

### A data-driven supervised learning approach identifies an association between PMC sulcal morphology and individual differences in face recognition ability

We next tested whether the sulci that show strong face selectivity (Figure 2, Figure 3) also show morphological relationships with face recognition ability. To do this, we implemented a supervised learning approach with the depth of each PMC sulcus predicting face recognition performance. Face recognition performance was measured using the Cambridge Face Memory Test (CFMT), a canonical face-identity recognition task.^50^ As in previous work, we ran a cross-validated LASSO regression—a data-driven feature selection technique—to identify sulci associated with CFMT performance (STAR Methods).^38,52–56^ This approach identifies the subset of features most strongly associated with the outcome while reducing overfitting, which is particularly important when the number of observations is modest relative to the number of predictors.^56–58^ As in prior work, to maximize the number of participants and sulci included in the model, we only included sulci that were present in the majority of hemispheres.^38^ Consequently, nine sulci were included in the LASSO regression. Eight were present in every hemisphere (ifrms, prcus-a, prcus-i, prcus-p, prculs-d, mcgs, spls, pos), while one sulcus (sspls-v) was present in a subset of hemispheres (Table S1), resulting in 35 NTs in the left-hemisphere model and 28 NTs in the right-hemisphere model. Given the modest ratio of observations (n) to predictors (K) in this analysis (n:K = 3.1 for the right-hemisphere model), the objective of this analysis was to establish stable associations between sulcal morphology and face recognition performance (i.e., feature selection), the predictive utility of which can be tested in future studies with larger samples.

A right-hemisphere LASSO regression predicting CFMT score from sulcal depth in NTs identified three PMC sulci associated with CFMT performance (ifrms_RH_: β_CV_ = -1.39, prcus-p_RH_: β_CV_ = -0.43, prcus-i_RH_: β_CV_ = -0.28; Figure 4a) at the α-value that minimized root mean squared error (RMSE), which was selected via fully nested leave-one-participant-out cross-validation (α = 1.35, RMSE_CV_ = 5.21; Figure 4b). The three sulci were consistently selected across cross-validation folds (ifrms_RH_ = 100%, prcus-p_RH_ = 89%, prcus-i_RH_ = 79%, all other sulci = <4%), suggesting adequate model stability. To formally test model stability, we bootstrapped this pipeline 2,000 times. Bootstrap validation showed that these sulci were selected in a majority of resamples (ifrms_RH_: 86%, prcus-p_RH_: 56%, prcus-i_RH_: 61%), with statistically significant coefficient estimates at *p*<.05 (95% CI: ifrms_RH_ [-4.41, -0.22], prcus-p_RH_ [-4.41, -0.03], prcus-i_RH_ [-5.27, -0.15]; see Table S2 for all sulci results). Critically, two of the three sulci identified by the model (prcus-p and prcus-i) are the same sulci that showed strong face selectivity in the fMRI analysis (Figure 2). To confirm that these effects were specific to the selected sulci and not a general effect of right-hemisphere PMC sulci, we used the corrected Akaike Information Criterion (AIC; see STAR Methods) to compare the selected model to a full model that included all nine sulci. The three-sulci LASSO model was preferred over the full model (*full model*: AIC = 151.0, *selected model*: AIC = 116.8; ΔAIC = 34.2). Despite the stability of the feature selection, the cross-validated adjusted R² for the selected model was low (R²_adj_ < .001). This indicates that while the feature selection identified stable feature–outcome associations between sulcal morphology and behavior, the relationships are modest relative to overall individual variability, and larger sample sizes would be needed for reliable out-of-sample prediction.

**Figure 4:**
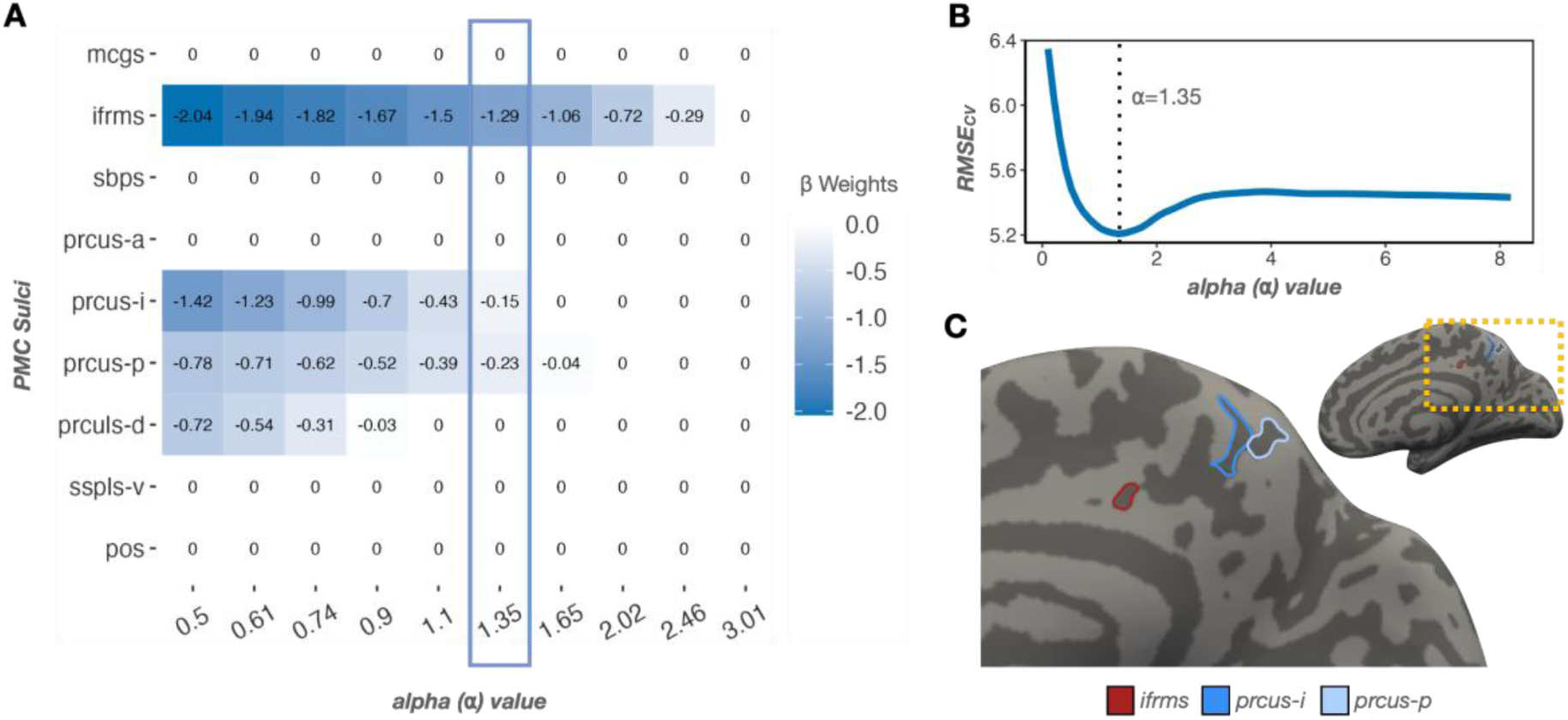
A data-driven supervised learning approach identifies an association between PMC sulcal morphology and individual differences in face recognition ability. (A) Top: We implemented a data-driven supervised learning approach to test if sulcal depth—a defining morphological feature of primary (deep), secondary (intermediate), and tertiary (shallow) sulci^59–75^—was associated with performance on a face recognition task (Cambridge Face Memory Test, CFMT). Standardized beta-coefficients for each sulcus at a range of shrinking parameter (α) values were estimated from a LASSO regression predicting CFMT score from sulcal depth in the right hemisphere of NTs. The highlighted box indicates coefficients at the optimal α-value, which was determined with fully-nested leave-one-participant-out cross-validation. The model identified three recently identified PMC sulci in the right hemisphere whose sulcal depth was associated with CFMT performance: the inframarginal sulcus (ifrms; β = -1.29), the intermediate precuneal sulcus (prcus-i; β = -0.15), and the posterior precuneal sulcus (prcus-p; β = -0.23). (B) Cross-validated root mean squared error (RMSE_CV_) for a wide range of α-values, with indication of the α that minimized RMSE_CV_ (dotted line). (C) Inflated cortical surface reconstruction of an example participant showing the location of the three right-hemisphere sulci selected by the LASSO regression.

Control analyses showed that effects were specific to sulcal depth in the right hemisphere: a left-hemisphere LASSO regression selected no sulci at the optimal α-value (α = 2.3, RMSE_CV_ = 5.49), and a right-hemisphere LASSO regression implemented with cortical thickness instead of depth selected no sulci at the optimal α-value (α = 4.49, RMSE_CV_ = 5.44). Further, these findings may be specific to neurotypical participants: implementing the same approach in DPs resulted in no sulci being selected in either hemisphere (right: α = 3.68, RMSE_CV_ = 4.29; left: α = 2.15, RMSE_CV_ = 4.08). However, the right-hemisphere results in DP participants should be interpreted with caution, as hemispheric differences in sspls-v incidence (see Figure 3) led to limited sample size for that model (lh: N=30, rh: N=16).

Together, these results identify stable associations between right PMC sulcal morphology and face recognition performance that can be further validated in future studies. Critically, two of the three identified sulci (prcus-p and prcus-i) from this structural–behavioral analysis also showed strong face selectivity as identified by fMRI (Figure 2), showing convergent evidence from both structural and functional analyses that these sulci are involved in face processing.

## Discussion

In the present study, we examined the relationship between sulci in posteromedial cortex (PMC) and face processing using structural, functional, and behavioral measures. We report three main findings. First, we found that recently defined sulci in posterior, but not anterior, PMC were face-selective as measured with fMRI, and this result was consistent across NTs and DPs. Second, we found that a recently identified sulcus in posterior–ventral PCC (sspls-v) was present less frequently and exhibited lower face selectivity in DPs compared to NTs in the right, but not left, hemisphere. Third, using a data-driven supervised learning approach, we found that the depths of three recently identified PMC sulci in the right, but not left, hemisphere were associated with performance on a face recognition task. In the sections below, we discuss these findings in the context of (1) sulcal-functional couplings in human face-selective regions beyond PMC, (2) situating the PMC face-selective region (“medial face area”) within the core and extended aspects of the human face processing network, and (3) a possible role for the sspls-v in processing face familiarity.

### Sulcal-functional coupling of human face-selective regions

Classic models of face processing identify coarse anatomical-functional relationships between face-selective regions and anatomical structures. For example, the fusiform gyrus (FG) and superior temporal sulcus (STS) are often associated with different aspects of face processing, with the STS being associated more with facial motion and expression than the FG.^9,16,82–83^ Nevertheless, the FG and STS are large structures spanning several centimeters and recent findings have pinpointed more precise landmarks that are associated with face-selective regions in the human cerebral cortex. For example, there is more than one face-selective region on the FG,^76^ and the antero-lateral tip of the mid-fusiform sulcus predicts the location of one of these regions.^61^ Interestingly, a recent fMRI study showed that seeding regions of interest in a similar location lateral to the MFS during a resting-state scan revealed a functionally connected region in the posterior aspect of PMC.^5^ Close examination of the figures from that study indicates that this functional region also overlaps with prcus-p and extends into prcus-i, a finding that is consistent with the sulcal-functional coupling identified in the present study. It is particularly intriguing given that this region exhibited preferential BOLD responses to recall of familiar people over familiar places (Figure S1f), suggesting selective processing for faces.

Previous research has also identified face-selective regions in different aspects of human lateral prefrontal,^84–90^ parietal,^87–89,91–94^ and temporal cortices.^87–91,95–96^ Given findings from the present study—in which our data-driven approach identified two precuneal sulci (prcus-i and prcus-p) that are face-selective and whose morphology relates to face processing ability—future studies can expand on this work by testing (1) if there are sulcal-functional relationships in the lateral prefrontal, parietal, and temporal cortices, and (2) if those sulcal-functional relationships are also related to face processing ability. Further, the present work indicates that the sulcal-functional coupling between the prcus-p and the face-selective region in PMC also exists in individuals with DP. Future work will indicate if this sulcal-functional coupling extends to other clinical populations that have difficulties with face processing, and if they show a similar anterior–posterior gradient of face selectivity in the precuneus.

### PMC: Part of the core or extended system of face processing?

Previous theories propose that face processing in PMC is part of the “extended” system of face processing that integrates additional information with the visual image of the face such as social and emotional cues as well as familiarity, person knowledge, and episodic memories.^9,10^ Indeed, the sulci associated with face recognition performance in the present study (prcus-p, prcus-i, ifrms) overlap with the parietal memory network^98^ and the posterior medial network^99^ supporting long-term memory. Further, the sulcus that differs structurally and functionally in DPs (sspls-v) overlaps with a default mode subnetwork supporting episodic memory.^100^ Conversely, regions associated with the “core” system of face processing are typically assumed to process the visual features of the face itself.^9,10^ While recent research shows that the cortical expanse around prcus-p and prcus-i preferentially processes familiar faces and people (Figure S1)^4–8,133^—suggesting a more “extended” process is at play—our findings indicate that those task manipulations are not necessary to elicit face-selective activations in the posterior PMC. The present study used a traditional functional localizer experiment with unfamiliar face stimuli, which nevertheless revealed a sulcal-functional gradient of face selectivity along an anterior–posterior axis in PMC. Therefore “extended” processing of familiarity or semantically related person knowledge do not explain the fMRI findings from the present study.

Indeed, several studies support a role for posterior PMC in the visual processing of faces that goes beyond familiarity processing. One recent fMRI study conducted a group analysis of functional localizer data (faces > all other categories) from 787 participants from the Human Connectome Project.^8^ Close examination of the group-average cortical surfaces from that study shows a face-selective region overlapping with prcus-p and extending into prcus-i that is consistent with the structural-functional coupling identified in individual participants in the present work (Figure S1b). These findings are also consistent with a resting-state functional connectivity analysis that identified a cross-species “visual posterior precuneal region” that is structurally and functionally dissociable in both humans and macaques,^34^ as well as a recent precision-fMRI study that identified a posterior precuneal region sensitive to semantic representations of visual stimuli.^101^ Examining figures from both of these studies shows that, while the ifrms and prcus-p both overlap with regions associated with recognition memory, the prcus-p, but not the ifrms, overlaps with a region associated with visual processing. This pattern is consistent with findings from the present study in which the ifrms and prcus-p were both morphologically associated with face recognition performance, but only the prcus-p was face-selective as measured by fMRI. Further research is needed to better understand these sulcal-functional couplings in PMC, including investigating their theoretical implications and possible genetic underpinnings.^6^ Future studies can also add network-level insights by relating PMC sulcal morphology to white matter architecture, as discussed in previous studies of sulcal morphology and cognition.^43,52–53,55,59,78,104–106^

### A role for the sspls-v in processing face familiarity?

Sulcal-functional couplings in the human cerebral cortex can only exist if the sulcus itself is present, which is not always the case. There are extensive individual differences in the morphology of sulci, especially the presence or absence of the smaller, shallower, and more variable tertiary sulci. Perhaps the most widely explored variable sulcus is the paracingulate sulcus (PCGS), the presence or absence of which affects the underlying cytoarchitecture of anterior cingulate cortex.^77,102^ PCGS morphology also has behavioral and clinical implications.^77,103–106^ For example, the length of the PCGS predicts whether individuals with schizophrenia will hallucinate or not.^104^ Similarly, our recent work showed that the presence or absence of a putative tertiary sulcus in lateral prefrontal cortex is related to reasoning performance in both children and adults.^80–81^

In the present study, we documented that the recently identified ventral sub-splenial sulcus (sspls-v) is present less often in DPs than NTs and, when present, has lower face selectivity in DPs than NTs. These findings complement three previous findings showing that this ventral portion of PMC is involved in the processing of familiar faces. Afzalian and Rajimehr^8^ showed that the ventral, but not dorsal, portion of posterior PMC showed preferential responses to familiar faces using fMRI, which Woolnough and colleagues^7^ confirmed with intracranial recordings (Figure S1). Additionally, Visconti di Oleggio Castello and colleagues^4^ used fMRI multivariate pattern analysis to show that distinct but overlapping regions in PMC hold face identity and familiarity information, respectively—face identity information in a posterior precuneal region and identity-independent familiarity information in a ventral-posterior PCC region. These findings are consistent with a meta-analysis showing that the cortical expanse around sspls-v is strongly associated with autobiographical memory (Figure S1c).^2^ As such, it may be that the presence or absence of the sspls-v affects a person’s ability to process familiar faces, a hypothesis that can be tested in future research. If so, this would support the intriguing possibility that people with DP have intact development of a face-selective visual-processing area in the posterior PrC, but impaired development of a face-selective familiarity-processing area in ventral PCC.

## Conclusion

The present work identifies a local sulcal network in PMC that supports face processing with a right-hemisphere dominance. This sulcal network contains a recently identified sulcus that structurally and functionally differs between DPs and NTs; to our knowledge, this is the first evidence of a structural neuroanatomical marker in PMC that distinguishes individuals with face processing deficits. By focusing on individual sulcal anatomy—especially recently uncovered, putative tertiary sulci^36–37^—the present study enables a more precise understanding of the neuroanatomical substrate of face processing in PMC, and how this substrate differs in individuals with DP. These findings will serve as a foundation for future studies investigating face processing in PMC in both neurotypical and clinical populations.

## Acknowledgements

This research was supported by an NSF CAREER Award 2042251 (K.S.W.), NSF GRFP Award DGE-2234667 (J.P.K), NIH T32 NS047987 (J.P.K.), and NIH MSTP T32 GM140935 (E.H.W.). Data collected in London was supported by an ESRC grant (RES-061-23-0400) to B.D.; data collection at Dartmouth was supported by a Rockefeller Foundation award. Young adult neuroimaging and behavioral data were provided by the WU-Minn HCP Consortium (Principal Investigators: David Van Essen and Kamil Ugurbil; NIH Grant 1U54-MH-091657) and funded by the 16 NIH Institutes and Centers that support the NIH Blueprint for Neuroscience Research, and the McDonnell Center for Systems Neuroscience at Washington University. We thank Willa Voorhies and Jacob Miller for their assistance developing the data analysis pipeline used for this study.

## Author Contributions

J.P.K., E.H.W., B.J.P., and K.S.W. designed research. X.C., G.J., L.G., Z.Z., and B.D. performed data collection and preprocessing. J.P.K., E.H.W., B.J.P., S.A.M., and K.S.W. performed research. J.P.K., E.H.W., and K.S.W. analyzed data. J.P.K. and K.S.W. wrote the paper. All authors edited the paper and gave final approval before submission.

## Declaration of Interests

The authors declare no competing interests.

## Star Methods

### Key resources table

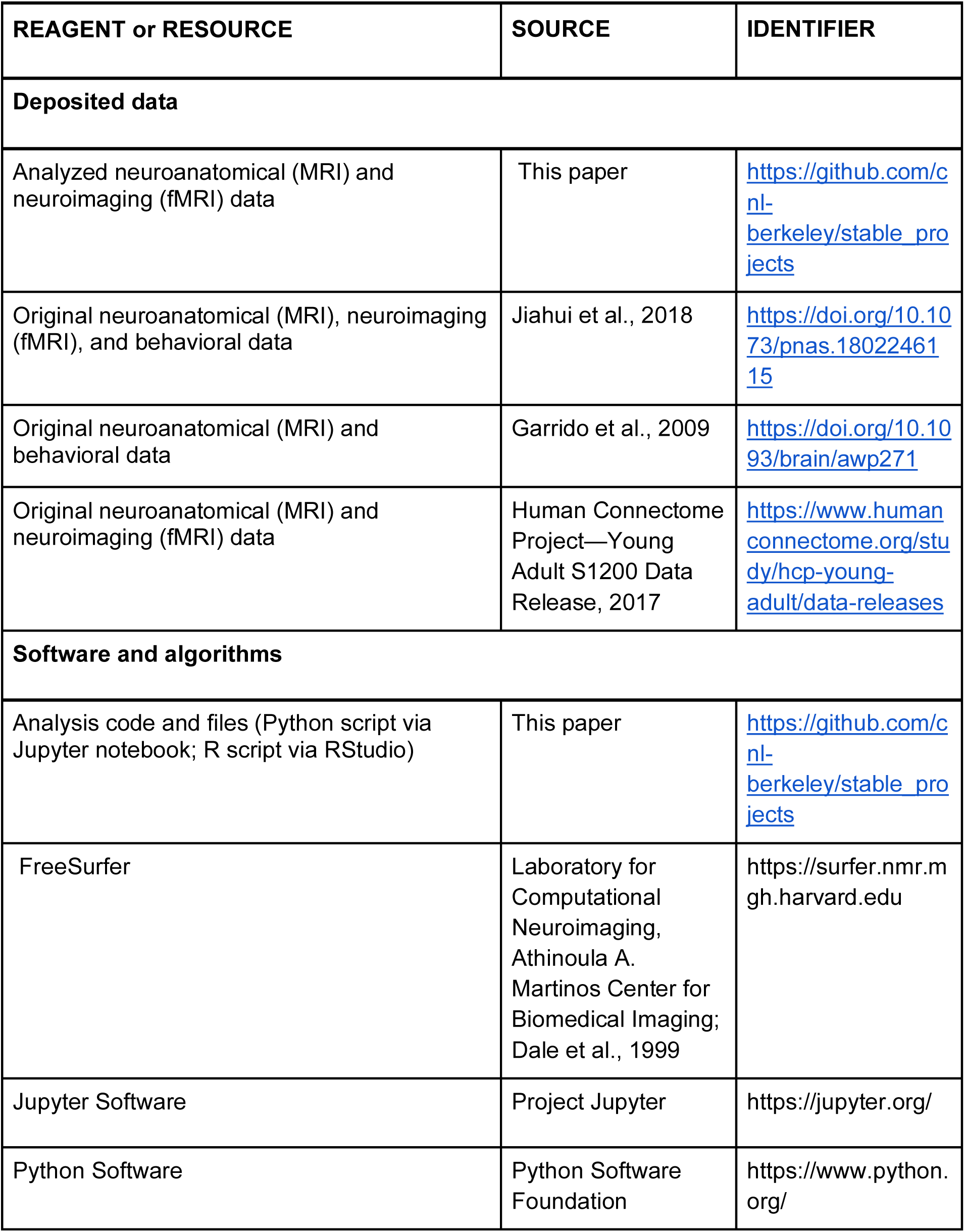
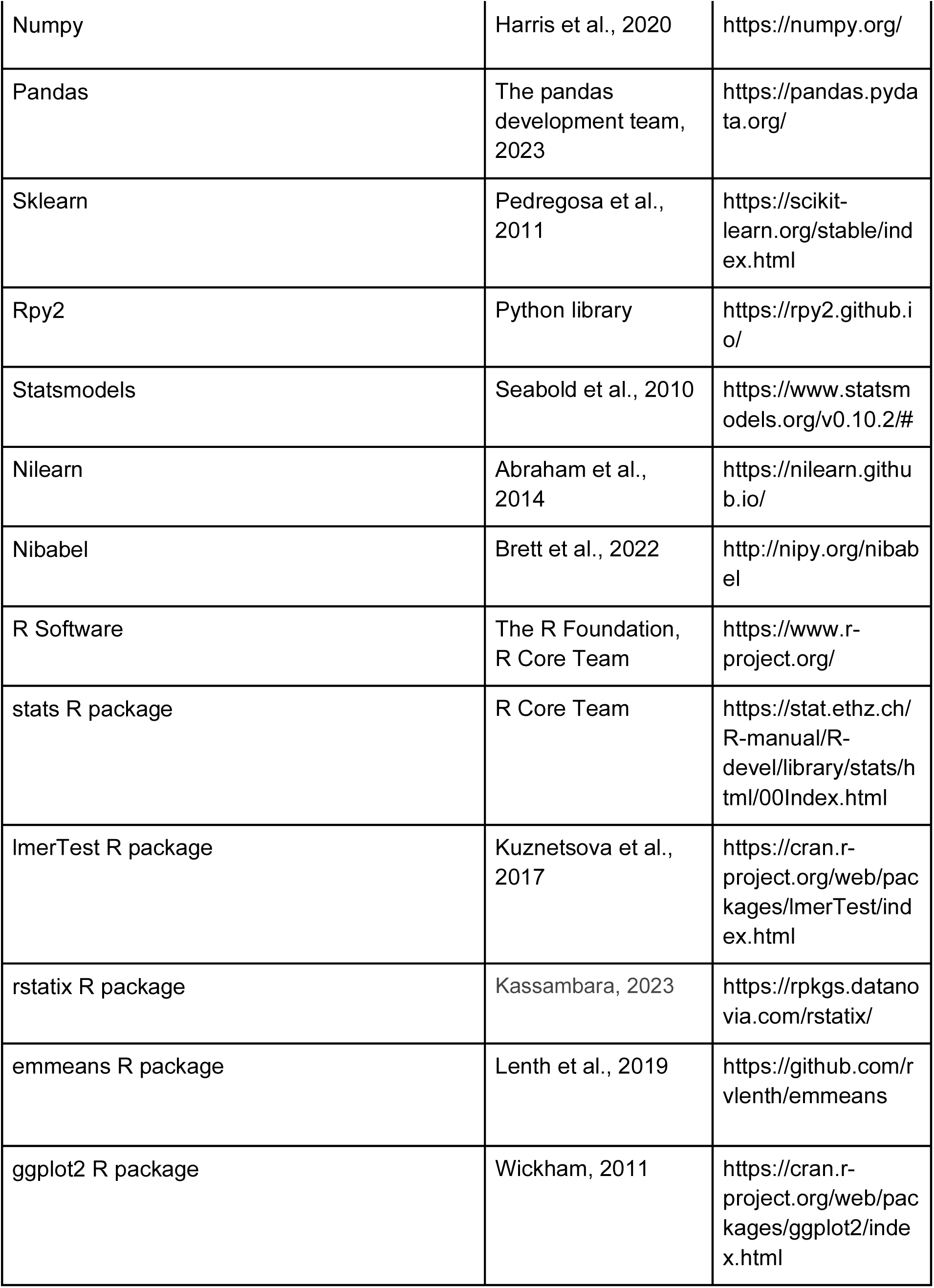
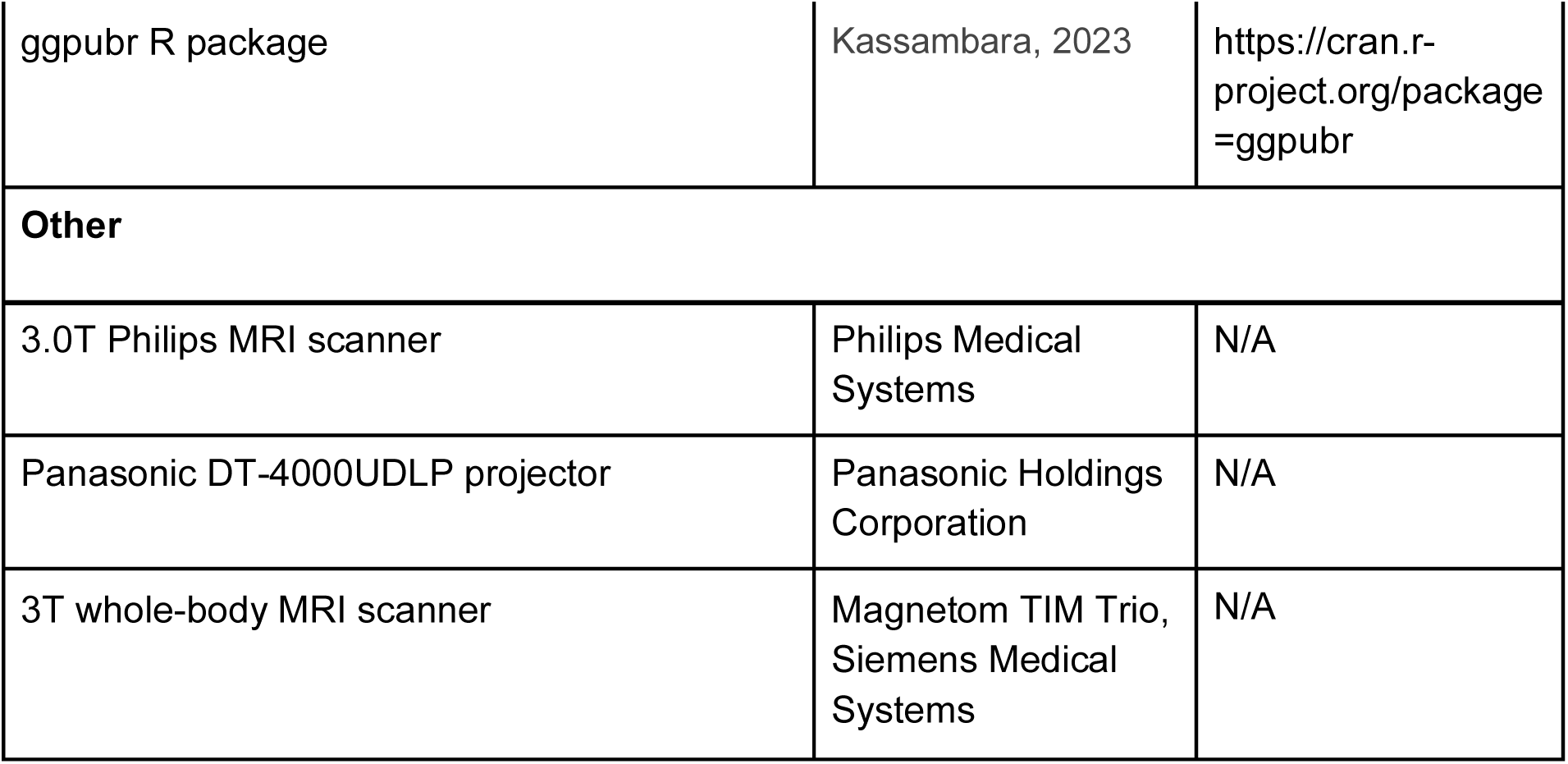

### Resource availability

#### Lead contact

- Further information and requests for additional resources should be directed to and will be fulfilled by the lead contact, Kevin Weiner.

#### Materials availability

- This study did not generate new unique reagents.

#### Data and code availability

- Processed data used for the present project have been deposited on GitHub and are publicly available as of the date of publication (see key resources table).
- All original code used for the present project has been deposited on Github and is publicly available as of the date of publication. DOIs/links are listed in the key resources table.
- Any additional information required to reanalyze the data reported in this paper is available from the lead contact upon request.

#### Experimental Model and Subject Details

Our analyses were conducted using three previously published datasets from Jiahui and colleagues^46^ (Dataset 1), Garrido and colleagues^40^ (Dataset 2), and the Human Connectome Project–Young Adult S1200 Data Release (Dataset 3; https://www.humanconnectome.org).^47,49^ Datasets 1 and 2 were used for the structural–behavioral analyses relating sulcal depth to face recognition scores, and sulcal incidence to group, while Datasets 1 and 3 were used to analyze the face selectivity of PMC sulci. Descriptive details of these datasets relevant to the present study are reproduced below.

#### Participants (Dataset 1)

Twenty-two participants with developmental prosopagnosia (DPs: 15 females, Age mean (sd) = 41.9 (12.0), range: 25–62) and 25 neurotypical controls (NTs: 15 females, Age mean (sd) = 42.3 (10.1), range: 21–55) participated in the study. DP participants were recruited from www.faceblind.org, and all reported problems in daily life with face recognition. To assess their face recognition, DPs were tested with the Cambridge Face Memory Test (CFMT),^50^ a famous faces test (FFT),^51^ and an old–new face discrimination test.^51^ All but one DP participant performed two or more standard deviations below the mean of published control results in at least two of the three diagnostic tests.^97^ The DP participant who did not reach -2 standard deviations on two tests scored poorly on two of the three tasks (CFMT: z = -1.9; FFT: z = -7.1; old–new: z = -0.5), so we included them to increase the sample size. All participants had normal or corrected -to-normal vision and had no current psychiatric disorders. Participants were leveraged from prior studies on category selectivity and ventral-temporal sulcal morphology in DP individuals.^31,46^

#### Participants (Dataset 2)

Seventeen individuals with DP (11 females, Age mean (sd) = 30.9 (7.5), range: 20–46) and 18 NTs (11 females, Age mean (sd) = 28.9 (5.7), range: 23–43) participated in the study. All reported being right-handed. All 35 participants showed normal or corrected to normal visual acuity when tested with Test Chart 2000 (Thompson Software Solutions, Hatfield, UK). DP participants contacted the Duchaine lab through http://www.faceblind.org and reported significant difficulties recognizing familiar faces in everyday life. To ascertain that the face recognition deficits that the participants were reporting were consistent with DP, each individual was tested on the CFMT and FFT.^50,51^ All DPs performed significantly below the mean of published NT means for these two tests. Structural MRI data from the same participants was reported in prior studies of voxel-based morphometry and ventral temporal sulcal morphology in DPs.^40,43^

#### Participants (Dataset 3)

Data for the Human Connectome Project neurotypical adult cohort (HCP NT) analyzed in the present study were sourced from the freely available HCP database (https://humanconnectome.org/study/hcp-young-adult).^47,49^ The dataset consisted of the first five numerically listed HCP participants and a random selection of 66 additional participants for a total of 71 participants (36 females, Age mean (sd) = 28.97 (3.78), range 22–36). These participants are the same that were used in previous studies characterizing the sulcal anatomy of PMC.^36–37^

## Method Details

### Face processing behavioral tasks (Datasets 1 and 2)

In the present study, we used the CFMT as our quantitative face processing measure for three main reasons. First, the CFMT is a commonly used measure of unfamiliar face recognition that strongly correlates with tests of face perception and long-term face memory.^50,109^ Second, the CFMT is a well-established test with high reliability: previous studies report a Spearman-Brown split-half reliability of .90, as well as a test-retest reliability of .70 and alternate forms of reliability of .76.^109^ Third, we used the CFMT to maximize the statistical power of the LASSO analysis. All participants in Datasets 1 and 2, except one NT, completed the CFMT (N=82).

### MRI data acquisition

#### Dataset 1

All participants were scanned in a 3.0T Philips MRI scanner (Philips Medical Systems, WA, USA) with a SENSE (SENSitivity Encoding) 32-channel head coil. A high-resolution anatomical volume was acquired at the beginning of the scan using a high-resolution 3D magnetization-prepared rapid gradient-echo sequence (220 slices, field of view = 240 mm, acquisition matrix = 256 x 256, voxel size = 1 x 0.94 x 0.94 mm).

#### Dataset 2

All participants were scanned on a 3T whole-body MRI scanner (Magnetom TIM Trio, Siemens Medical Systems, Erlangen, Germany) operated with a radio frequency body transmit and 12-channel receive head coil. For each participant, a T1-weighted (T1w), 3D-modified driven equilibrium Fourier transform (MDEFT) dataset was acquired in sagittal orientation with 1 mm isotropic resolution (176 partitions, field of view = 256 x 240 mm^2^, matrix 256 x 240 x 176) with the following parameters: repetition time = 7.92 ms, echo time = 2.48 ms, inversion time = 910 ms (symmetrically distributed around the inversion pulse; quot = 50%), flip angle α = 16x, fat saturation, bandwidth 195 Hz/pixel. The sequence was specifically optimized for reduced sensitivity to motion, susceptibility artifacts, and B1 field inhomogeneities.

#### Dataset 3

Anatomical T1-weighted MRI scans (0.7-mm voxel resolution) were obtained in native space from the HCP database, along with outputs from the HCP-modified FreeSurfer pipeline (v5.3.0).^110–113^ Additional details on image acquisition parameters and image processing can be found in work by Glasser and colleagues.^110^ All subsequent sulcal labeling and extraction of anatomical metrics were calculated from the cortical surface reconstructions of individual participants generated through the HCP’s custom-modified version of the FreeSurfer pipeline.^110^

### Functional localizers

#### Dataset 1^46^

A detailed description of the functional scanning parameters has been previously described by Jiahui and colleagues.^46^ Here, we provide a brief overview. Participants completed a one-back task during a dynamic localizer scan containing five visual categories (faces, scenes, bodies, objects, and scrambled objects). Each participant completed five scans, composed of 10 12-s category blocks of video clips interleaved with 12-s fixation blocks (4.2 minutes in total). Each visual category was displayed twice in each scan in a quasi-random order. In each category block, six 1,500-ms video clips were presented interleaved by blank fixation screens presented for 500 ms. Stimuli were presented using SuperLab 4.5.3 (https://cedrus.com/superlab/index.htm) and displayed to the participant via a Panasonic DT-4000UDLP projector (resolution: 1,024 × 768; refresh rate: 60 Hz) at the rear of the scanner.

The five runs were then divided into localization runs and test runs to carry out a “leave-one-out” analysis.^114–115^ In each of the leave-one-out combinations, four of the five runs for a participant were used to localize the voxels that showed the strongest preference for the preferred category. To avoid the double-dipping problem,^116^ the responses of the selected voxels to each stimulus condition were then measured in the left-out run. All five combinations were analyzed and then averaged to produce the final result for each participant. Finally, face-selectivity maps were created from the difference between the response to faces vs. the response to objects for each participant.

#### Dataset 3

Face-selective regions in HCP NT participants were localized using a task in which four stimulus types (faces, places, tools, and body parts) were presented in separate blocks.^47^ The localizer consisted of two runs, and each run contained eight task blocks (10 trials of 2.5 s each, for 25 s) and 4 fixation blocks (15 s each). Within each run, half of the task blocks used a 2-back working memory task and the other half implemented a 0-back working memory task. A 2.5 s cue indicated the task type at the start of the block. For each trial, the stimulus was presented for 2 s, followed by a 500 ms inter-trial interval (ITI). Linear contrasts were computed to estimate effects of interest (e.g., faces vs. other categories). Fixed-effects analyses were conducted to estimate the average effects across runs within each participant.

### Cortical surface reconstruction

Cortical surface reconstructions for every participant from all three datasets were generated from their T1 scans using the standard FreeSurfer pipeline (v6.0.0; https://surfer.nmr.mgh.harvard.edu;).^111–113^ Cortical reconstructions were created from the resulting boundary made by segmenting the gray and white matter in each anatomical volume with FreeSurfer’s automated segmentation tools.^111^ After manual labeling of sulci, morphological metrics were extracted from individual participants’ cortical surface reconstructions using Freesurfer tools.

### Macroanatomical boundaries of the PCC and PrC within PMC

Based on the most recent and comprehensive atlas of sulcal definitions throughout the cerebral cortex^117^ as well as recent work identifying novel shallow sulci in posteromedial cortex (PMC),^36–37^ we define 12 sulci that border or are located within the posterior cingulate cortex (PCC) and the precuneus (PrC). Marking the anterior, posterior, and inferior boundaries, respectively, of PrC are the marginal ramus of the cingulate sulcus (mcgs), parieto-occipital sulcus (pos), and splenial sulcus (spls, also known as the subparietal sulcus or sbps).^117^ Contained within PrC are five sulci: three precuneal sulci known as the posterior precuneal sulcus (prcus-p), the intermediate precuneal sulcus (prcus-i), and the anterior precuneal sulcus (prcus-a), the dorsal precuneal limiting sulcus (prculs-d), and a recently identified shallow sulcus called the ventral precuneal limiting sulcus (prculs-v), which is not present in every hemisphere (see Table S1 for incidence rates of variably present sulci).^36–37^ PCC is bounded inferiorly by the callosal sulcus (cas), superiorly by the cingulate sulcus (cgs), mcgs, and spls, and posteriorly by the pos. Contained within PCC, and running its length along an anterior-dorsal to posterior-ventral axis, are four shallow sulci (PCC can be described as a tertiary “plane” or “corridor”): the posterior intracingulate sulcus (icgs-p), the inframarginal sulcus (ifrms), the dorsal sub-splenial sulcus (sspls-d), and the ventral sub-splenial sulcus (sspls-v).^36–37^ Of these putative tertiary sulci, the ifrms is present in every hemisphere, while the latter three are not.^36–37^

### Sulcal labeling

Each PCC and PrC sulcus was manually defined within each individual hemisphere on the *FreeSurfer* inflated mesh with tools in *tksurfer* as described in our previous work.^53,59^ Specifically, the *curvature* metric in FreeSurfer distinguished the boundaries between sulcal and gyral components, and manual lines were drawn to separate sulcal components. In some cases, the precise start or end point of a sulcus can be difficult to determine on one surface.^118^ Thus, we used the *inflated*, pial, and *smoothwm* surfaces of each individual to inform our labeling, form a consensus across surfaces, and clearly determine each sulcal boundary. The sulcal labels were generated using a two-tiered procedure. The labels were first defined manually by trained raters (J.K., E.W., and B.P.) and then finalized by a neuroanatomist (K.S.W.). In each hemisphere, we first labeled the deeper and more stable sulci bounding the PCC and PrC (e.g., pos, spls, mcgs), and then we identified the remaining sulcal components within the PCC and PrC. All anatomical labels for a given hemisphere were defined blind to participant group (i.e., NT vs. DP) and before any morphological or functional analyses were performed.

### Extracting incidence rates and anatomical features from sulcal labels

After all sulci were defined, incidence rates were calculated as the percentage of hemispheres in which a sulcus was present. Anatomical features [sulcal depth (standard Freesurfer units)] and mean cortical thickness (mm) were then extracted. Raw depth metrics were computed in native space from the .sulc file generated in FreeSurfer v6.0.0 using custom Python code leveraging various functions from the numpy, pandas, nilearn, and nibabel packages.^59,111^ Briefly, depth values are calculated based on how far removed a vertex is from what is referred to as a “mid-surface,” which is determined computationally so that the mean of the displacements around this “mid-surface” is zero. Thus, generally, gyri have negative values, while sulci have positive values. Given the shallowness and variability in the depth of tertiary sulci,^36–38,52–55,107–108^ some mean depth values extend below zero. We emphasize that this merely reflects the metric implemented in FreeSurfer. As in prior work,^36–38,43,52–55,107–108^ we normalized sulcal depth to the maximum depth value within each individual hemisphere in order to account for differences in brain size across individuals and hemispheres. All depth analyses were conducted using normalized mean sulcal depth. Mean cortical thickness (mm) was extracted from each sulcus using the built-in *mris_anatomical_stats* function in FreeSurfer.^119^

### Distinctions among primary, secondary, and tertiary sulci

As described in classic studies, as well as our previous work, tertiary sulci are defined as the last sulci to emerge in gestation after the larger and deeper primary and secondary sulci.^59–75,117^ While there is variability among individuals, previous studies suggest that, a) primary sulci emerge prior to 32 weeks in gestation, b) secondary sulci emerge between 32-36 weeks in gestation, and c) tertiary sulci emerge during and after 36 weeks.^64–65,67^ Tertiary sulci continue to develop postnatally.^64–68,139^

Based on this research and recent work defining putative tertiary sulci in PMC, we identified the mcgs, pos, and spls as primary sulci; the three precuneal sulci (prcus-p, prcus-i, prcus-a) and the prculs-d as secondary sulci; and the ifrms, sspls-d, sspls-v, and icgs-p as putative tertiary sulci.^36–37,102^ Defining these latter four sulci as tertiary sulci is further supported by their shallow depth and small surface area, which are defining morphological features of tertiary sulci. However, because the definition of primary, secondary, and tertiary sulci remains debated, we refer to tertiary sulci explored in the present study as putative. Future research leveraging noninvasive fetal imaging should further improve the distinctions among primary, secondary, and tertiary sulci. Critically, the data-driven approach adopted for our behavioral analyses (“Behavioral analysis” below) is agnostic to these distinctions; it quantitatively determines which sulci are most associated with face recognition performance regardless of their classification.

### Behavioral analysis

We used a data-driven supervised learning approach to determine which PMC sulci demonstrated a structural-behavioral relationship between sulcal depth and CFMT score. Our sample for behavioral analyses included 42 neurotypical adults [26 females, Age mean (sd) = 36.55 (10.85)], after removing one NT participant who did not have a CFMT score. To maximize the number of participants and sulci included in the model, we only included sulci that were present in the majority of hemispheres.^38^ Consequently, nine sulci were included in the LASSO regression. Eight were present in every hemisphere (ifrms, prcus-a, prcus-i, prcus-p, prculs-d, mcgs, spls, pos), while one sulcus (sspls-v) was present in a subset of hemispheres (see Table S1), resulting in 35 NTs in the left-hemisphere model and 28 NTs in the right-hemisphere model

#### Testing for associations between sulcal morphology and face processing performance

To assess the relationship between normalized sulcal depth and face processing performance in a way that maximized the rigor and interpretability of our findings, we implemented a two-pronged approach adapted from our previous work^38,52–55^ and current recommendations^56–58,131–132^.

1. *Regularization*: We used L1-regularization (LASSO regression) to select the optimal model. Regularization techniques provide a data-driven method for model selection and are recommended to improve the generalizability of a model.^56,131–132^ Unlike many techniques that only *assess* generalizability, L1 regularization increases generalizability by providing a sparse solution that reduces coefficient values and decreases variance in the model, thus guarding against overfitting without increasing bias.^56,132^
2. *Fully nested cross-validation:* In addition to using regularization techniques, our LASSO model was tuned with cross-validation, a within-sample method for improving the generalizability of a model that is a standard practice in machine learning.^57–58,131^ Cross-validation creates a series of within-sample train/test splits that allows for every data point to be used for both training and testing. In the present study, we used leave-one-participant-out cross-validation (LOOCV), in which the model is successively fit to every data point but one and then tested on the left-out data point. To further increase generalizability, cross-validation in this study was fully nested, meaning that parameter tuning and feature selection were performed separately for each cross-validation fold (i.e., all steps of model selection are fully “nested” within an outer cross-validation loop). Fully nested cross-validation is the most rigorous, least biased method for within-sample development of generalizable models.^57–58^

#### Model selection

We applied a least absolute shrinkage and selection operator (LASSO) regression model to determine which sulci, if any, were associated with CFMT performance. The depth of the nine consistently present PMC sulci were included as predictors in the regression model. LASSO performs L1-regularization by applying a penalty, or shrinking parameter (α), to the absolute magnitude of the coefficients such that:

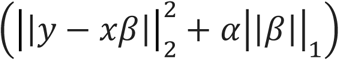

In a LASSO regression, low coefficients are set to zero and eliminated from the model. In this way, LASSO can facilitate variable selection, leading to simplified models with increased interpretability and reduced overfitting.^56,132^ By convention, we used cross-validation to select the shrinking parameter (α). We used the SciKit-learn *GridSearchCV* package^134^ to perform an exhaustive grid search across a wide range of α-values (0.1-10.0) and selected the value that minimized cross-validated root mean squared error (RMSE_CV_). Sulci chosen for the final model had non-zero coefficients at the optimal alpha value.

#### Model stability and validation

To assess the stability of feature selection, we performed bootstrap validation with 2,000 iterations. In each bootstrap iteration, we resampled the data with replacement and repeated the full LASSO pipeline, recording which features were selected and their coefficient values. We calculated 95% confidence intervals for each coefficient to test whether the associations were statistically significant (i.e., confidence intervals excluding zero).

#### Model comparison testing

To formally compare the LASSO-selected model with the full model in terms of estimated prediction error, we used the Akaike Information Criterion corrected for sample size (AIC). AIC provides an estimate of in-sample prediction error and is appropriate for model comparisons in which the number of parameters (*K*) in the largest model is high relative to the sample size (*n*) such that 𝑛/𝐾 ≤ 40.^136^ In the context of linear regression, the corrected AIC is given by:

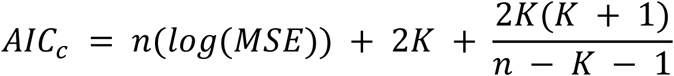

where *n* is the sample size and MSE is the mean squared error for the model.^138^ By comparing AIC scores, we assess the relative performance of the two models. If the Δ*AIC* is greater than 2, it suggests an interpretable difference between models. If the Δ*AIC* is greater than 10, it suggests a strong difference between models. The lower Δ*AIC* value indicates the preferred model.^135–136^ Lower Δ*AIC* values indicate better models that balance goodness-of-fit with model complexity.

#### Assessment of predictive performance

To evaluate the absolute predictive performance of the selected model, we calculated the cross-validated adjusted R^2^, which indicates how well the model predicts new observations relative to simply using the sample mean. This metric provides an assessment of whether the model achieves predictive utility, beyond any feature–outcome associations identified during model selection.

#### Control analyses

We conducted several control analyses to test the specificity of our findings: (1) We tested whether similar associations existed in the left hemisphere; (2) We examined whether cortical thickness (rather than sulcal depth) showed similar associations; and (3) We tested whether the same sulcal features predicted face recognition performance in participants with developmental prosopagnosia (DP). These control analyses help determine whether the observed sulcal– behavioral associations are specific to right-hemisphere sulcal depth in neurotypical participants.

### Functional analysis

Mean face selectivity was measured as the difference between mean activation for faces vs. other categories during a functional localizer task. In Dataset 1 (NTs and DPs), mean face selectivity was quantified as the percentage change in response to faces minus the percentage change in response to objects (% Faces - % Objects) averaged across the entire sulcus (as in Jiahui and colleagues^46^; see “Functional localizers”). In Dataset 3 (HCP NTs), mean face selectivity was quantified as the z-scored contrast of faces > all other categories (places, tools, body parts) averaged across the entire sulcus (as in Chen and colleagues^76^). The face selectivity of each sulcus was tested using mixed-model ANOVAs on mean face selectivity with fixed effects of sulcus, hemisphere, and group (except in the HCP NT analysis, in which there was no group comparison), and participant as a random effect. Hemisphere was nested within participant, and all interaction terms were also included in the model. Significant mean face selectivity against zero was tested using two-sided post hoc t-tests with Tukey’s adjustment for multiple comparisons. Hemispheric, group, and between-sulci differences in mean face selectivity were tested using the same model, again with Tukey’s adjustment for multiple comparisons.

### Quantification and Statistical Analysis

The structural-behavioral analyses relating sulcal depth to face processing scores were done in Python (v3.10.4; https://www.python.org/) and R (v4.2.3; https://www.r-project.org/) via JupyterLab notebook (v6.4.12; https://jupyter.org/) and the *rpy2* package, which enables R code to be run in Python environments. We used the Python *sklearn* library for the LASSO procedure, grid search, and cross-validation, and we used the Python *statsmodels* library for multiple regressions and model comparison F-tests. We used the *ggplot2* package (via *rpy2*) to generate data visualizations, and the *stat_cor* package to generate Spearman correlations. All other analyses, including analyses of sulcal incidence and face selectivity, were performed in R via RStudio (v2023.03.0+386; https://posit.co/download/rstudio-desktop/). Tests for hemispheric and group differences in sulcal incidence were performed with the *chisq.test* function. Mixed-model ANOVAs were performed using the *lmerTest* package for the linear mixed models, the *anova_test* package from the *rstatix* library for ANOVAs, and the *emmeans* package for post-hoc t-tests.

## Notes

### Competing Interest Statement

The authors have declared no competing interest.

