## Supplemental Materials for "Overlooked neuroanatomical markers of face processing and developmental prosopagnosia in posteromedial cortex"

### Supplement

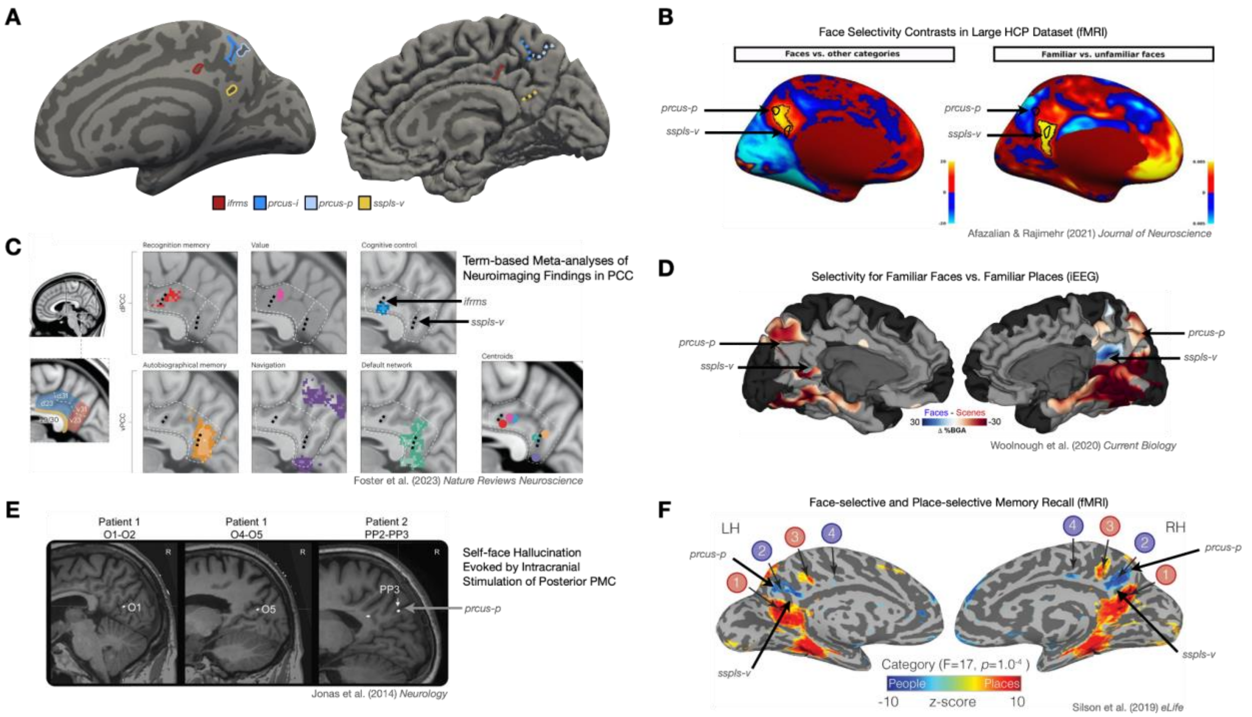


**Figure S1: Integrating findings from the present study with recent literature on the functional neuroanatomy of PMC** (A) Inflated surface (left) and pial surface (right) from an example participant from the present study. Four recently identified sulci (see legend) that we found to be significantly involved in face processing are labeled for ease of comparison with prior studies—two in the precuneus (prcus-p: posterior precuneal sulcus, prcus-i: intermediate precuneal sulcus) and two in the posterior cingulate cortex (sspls-v: ventral sub-splenial sulcus, ifrms: inframarginal sulcus).

(B) Results from Afzalian and Rajimehr[1] showing face selectivity contrasts (left: Faces vs. other categories; right: Familiar vs. unfamiliar faces) from a large sample of participants from the Human Connectome Project (HCP, N=787). Face selectivity (left, z-score) and familiar face selectivity (right, $p < .05$, FDR-corrected) are shown in hot colors, and the top 1% of selective voxels are outlined (black border). Outlines showing the approximate locations of two recently identified sulci (prcus-p, sspls-v) in posterior PMC are added (arrows) onto the group-average surface to facilitate comparison with the present study. The prcus-p and part of the sspls-v co-localize with a face-selective region in posterior PMC (left), while the sspls-v, but not the prcus-p, co-localizes with a region selective for familiar faces in ventral PCC (right).

(C) Results from Foster and colleagues[2] summarizing functional activations in PCC using term-based meta-analyses of neuroimaging studies. Anatomical scans (left panel) show cytoarchitectonic divisions within PCC, which correspond with functional dissociations across various types of cognitive tasks (center panels). Centroids summarizing the results from each meta-analytic search term are shown in the right panel. Labels (black dotted lines) showing the approximate locations of the ifrms and sspls-v on the average surfaces are added to facilitate comparison with the present study. The ifrms is located in a region associated with recognition memory and cognitive control, while the sspls-v is located in a region associated with autobiographical memory and the default mode network.

(D) Results from Woolnough and colleagues[3] showing regions selective for familiar faces (blue) vs. familiar places (red) as measured with intracranial EEG recordings (N=66). Approximate locations of the prcus-p and sspls-v are added (arrows) to the group-average surface to facilitate comparison with the present study. The sspls-v in the right hemisphere, but not the left, is located in a region that is strongly selective for familiar faces, which matches results from the present study.

(E) Results from Jonas and colleagues[4] showing the anatomical location of stimulation sites in the right-hemisphere posterior PMC of two epilepsy patients. Stimulation of these sites evoked autoscopic hallucinations in which patients saw their own face in the contralateral hemifield. The stimulation sites for Patient 1 were located in the parieto-occipital sulcus, while the stimulation site for Patient 2 was located in the prcus-p (label added to facilitate comparison with the present study).

(F) Results from Silson and colleagues[5] showing regions selective for people (blue) vs. places (red) as measured with fMRI in neurotypical participants (N=24) during a memory recall task. Results show an alternating pattern of memory recall between person- and place-selective regions along the ventral/posterior–dorsal/anterior axis, with the strongest activations in posterior PMC (ROIs 1 and 2; numbered ROIs are from the original figure). Labels showing the approximate locations of the prcus-p and sspls-v are added (long arrows) to the group-average surface to facilitate comparison with the present study.

**Face selectivity of all sulci in PMC**

As stated in the main text, we tested for face selectivity in PMC sulci using data from a block-design fMRI experiment containing images of faces, scenes, objects, and bodies.[6] A mixed-model ANOVA for mean face selectivity [faces - objects; % signal change (as in Jiahui and colleagues)[6]] with fixed effects of sulcus, hemisphere (left, right), and group (NT, DP) identified a three-way interaction [$F(11,414) = 2.80$; $p = .002$; $\eta^2_G = .017$]. Post hoc analyses revealed several additional findings. Significant face selectivity was found in the right hemisphere of NTs in the sspls-v, prcus-i, prcus-p, prculs-v, prculs-d, and spls ($p$s ≤ .037, all one-sample t-tests Bonferroni-corrected for multiple comparisons), with the strongest selectivity in a subset of sulci located in posterior PCC (sspls-v: $p < .0001$) and posterior PrC (prcus-p: $p < .0001$; prculs-v: $p < .0001$). In the left hemisphere of NTs, significant face selectivity was found in posterior PrC in the prcus-i and prcus-p ($p$s < .0001) as well as the prculs-v ($p = .022$). In DPs, significant face selectivity was found in the right hemisphere in the prcus-i, prcus-p, prculs-v, prculs-d, and spls ($p$s ≤ .008), with the strongest selectivity in posterior PrC (prcus-p: $p < .0001$). In the left hemisphere of DPs, the prcus-i, prcus-p, prculs-v, spls, and sspls-d showed significant face selectivity ($p$s ≤ .011), with the strongest selectivity again found in posterior PrC (prcus-p: $p < .0001$). Across both hemispheres and in both groups, the only sulcus selective for objects over faces was the pos ($p$s ≤ .011). As stated in the main text, we ran the same analysis in a sample of 71 participants from the Human Connectome Project (HCP) to assess whether similar results were present in an independent dataset. A mixed-model ANOVA for mean face selectivity [faces - all other categories; z-score (as in Chen and colleagues)[76]] with the same fixed effects as above yielded an interaction effect between sulcus and hemisphere [$F(11,649) = 4.46$; $p < .0001$; $\eta^2_G = .021$]. Post hoc analyses revealed significant face selectivity in the sspls-v, prcus-p, prculs-v, and spls ($p$s ≤ .003), with the strongest selectivity in posterior PCC (sspls-v: $p$ < .0001) and posterior PrC (prcus-p: $p$ < .0001; prculs-v: $p$ < .0001). In the left hemisphere of HCP NTs, significant face selectivity was found in the prcus-i, prcus-p, prculs-v, spls, and sspls-d ($p$s ≤ .008), with the strongest selectivity in posterior PrC (prcus-p: $p$ < .0001, prculs-v: $p$ < .0001). The only sulci showing a negative effect—i.e., selective for all other categories over faces—were the pos ($p$s ≤ .0001) and the prculs-d ($p$s ≤ .006).

| | **Neurotypical (NT, N=43)** | | **Dev. Prosopagnosia (DP, N=39)** | |
|---|---|---|---|---|
| | **Left (%)** | **Right (%)** | **Left (%)** | **Right (%)** |
| ***icgs-p*** | 20 (47) | 23 (53) | 19 (49) | 20 (51) |
| ***ifrms*** | 43 (100) | 43 (100) | 39 (100) | 39 (100) |
| ***sspls-d*** | 18 (42) | 21 (49) | 16 (41) | 16 (41) |
| ***sspls-v*** | 36 (84) | 29 (67) | 30 (77) | 16 (41) |
| ***prculs-v*** | 15 (35) | 17 (40) | 16 (41) | 18 (46) |

**Table S1: Incidence of Putative Tertiary Sulci in PMC**
Frequency counts and percentages by group (NT, DP) and hemisphere (Left, Right) for each putative tertiary sulcus in PMC. All other sulci in PMC were present in every hemisphere.

| | **Hemisphere** | **Proportion of iterations selected (%)** | **Median Coefficient** | **95% CI Lower Bound** | **95% CI Upper Bound** |
|---|---|---|---|---|---|
| ***prcus-i*** | RH | 61.3 | -1.85 | -5.27 | -0.15 |
| ***ifrms*** | RH | 86.5 | -1.76 | -4.41 | -0.22 |
| ***prcus-p*** | RH | 56.2 | -1.26 | -4.41 | -0.03 |
| ***prculs-d*** | RH | 53.4 | -1.23 | -3.89 | 1.86 |
| ***prcus-a*** | RH | 34.9 | -0.78 | -4.00 | 2.45 |
| ***pos*** | RH | 48.6 | -0.51 | -2.91 | 2.26 |
| ***sbps*** | RH | 41.3 | -0.47 | -3.14 | 3.11 |
| ***sspls-v*** | RH | 43.4 | 0.04 | -1.97 | 2.86 |
| ***mcgs*** | RH | 34.2 | 0.14 | -2.95 | 1.89 |

**Table S2: Model Stability Analysis of Right-hemisphere LASSO Regression**
Coefficient statistics for each sulcus after bootstrapping ($n_{iterations}$=2,000) the right-hemisphere LASSO regression in NT individuals. Sorted by median coefficient value. The three sulci selected by the original LASSO model were statistically significant at $p$<.05.

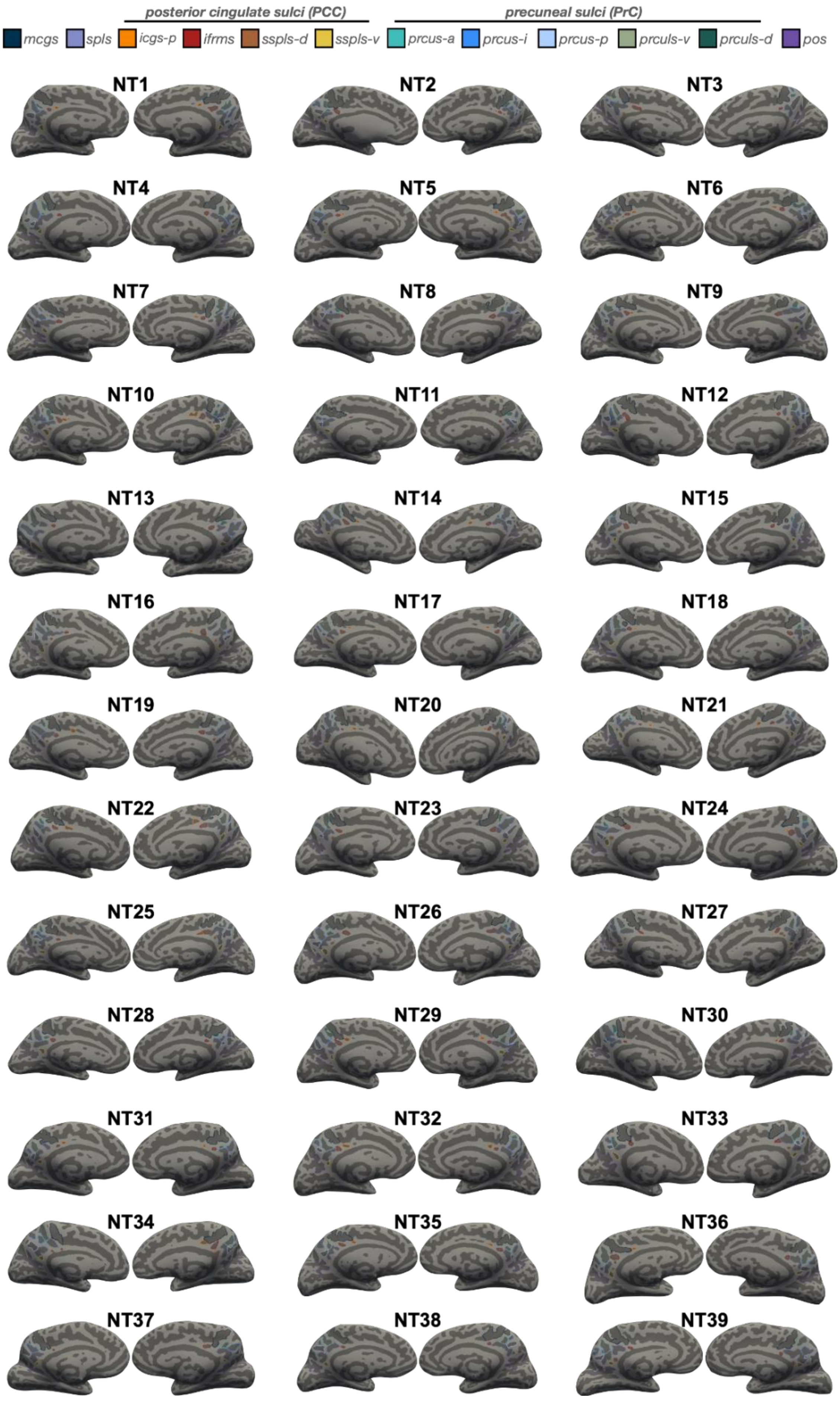
posterior cingulate sulci (PCC)
precuneal sulci (PrC)
mcgs
spls
icgs-p
ifrms
sspls-d
sspls-v
prcus-a
prcus-i
prcus-p
prculs-v
prculs-d
pos
NT1
NT2
NT3
NT4
NT5
NT6
NT7
NT8
NT9
NT10
NT11
NT12
NT13
NT14
NT15
NT16
NT17
NT18
NT19
NT20
NT21
NT22
NT23
NT24
NT25
NT26
NT27
NT28
NT29
NT30
NT31
NT32
NT33
NT34
NT35
NT36
NT37
NT38
NT39

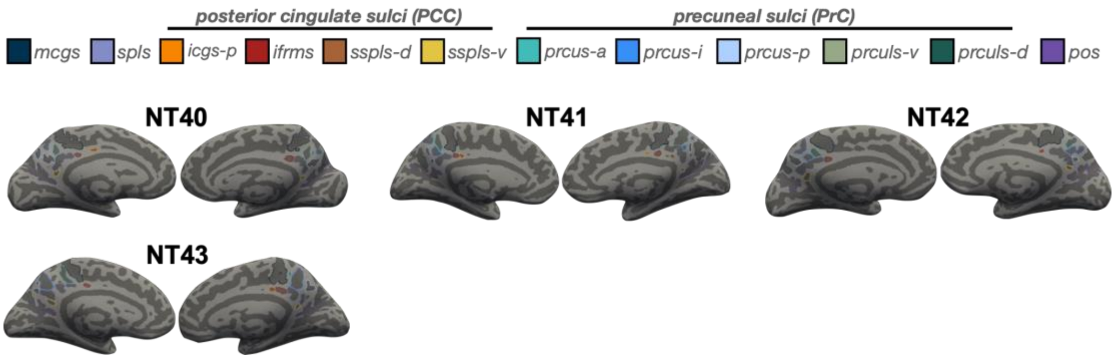


**Figure S2a. Manual PMC sulcal labels for every neurotypical (NT) participant**
Each sulcus is displayed on the left and right hemisphere inflated cortical surfaces in FreeSurfer 6.0.0, with each label displayed as an outline according to the key at the top. Each hemisphere contains at least 8 sulci (from anterior to posterior): ifrms, mcgs, spls, prcus-a, prcus-i, prcus-p, prculs-d, pos. An additional 4 sulci are variably present: icgs-p, sspls-d, sspls-v, and prculs-v.

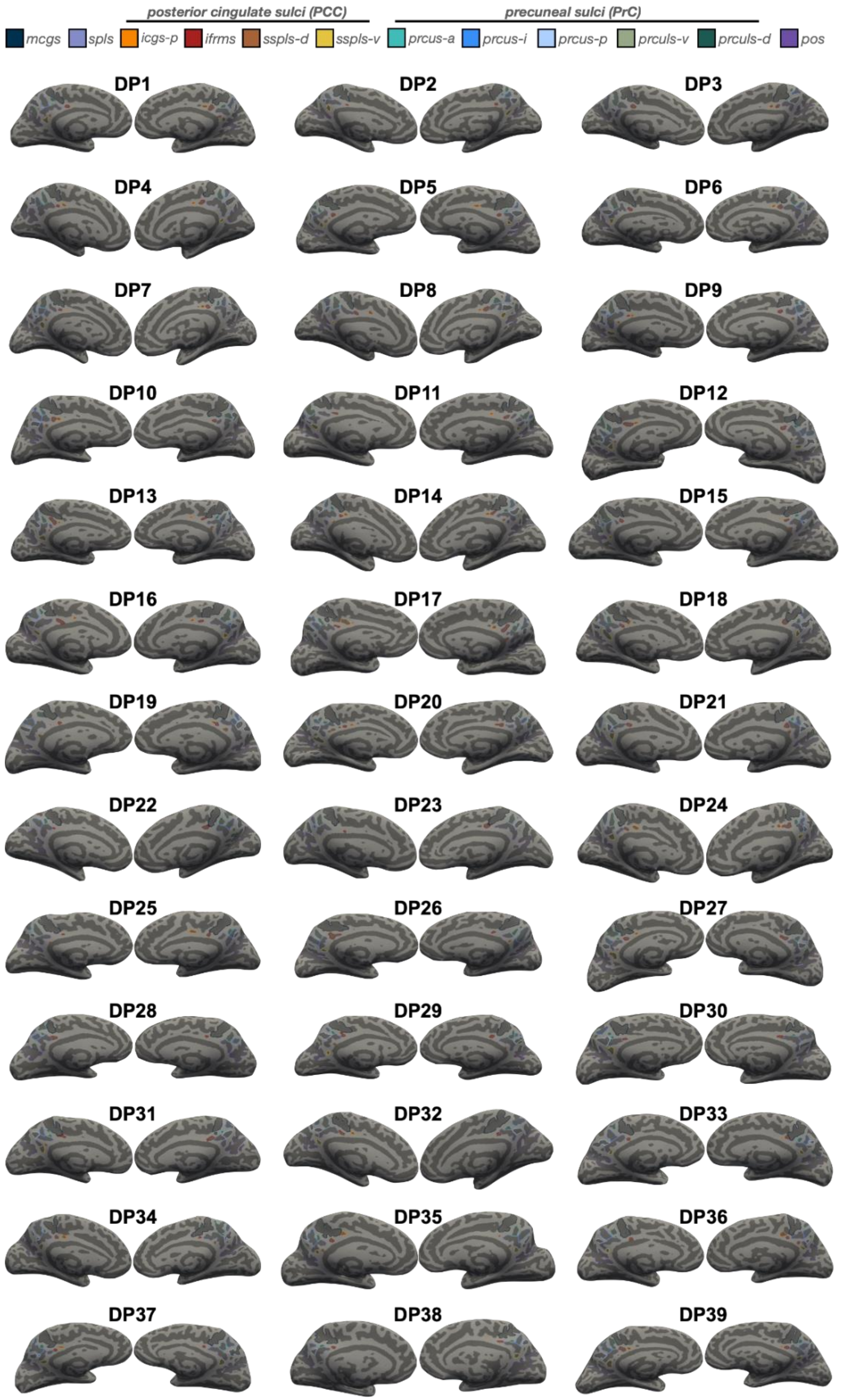

posterior cingulate sulci (PCC)
precuneal sulci (PrC)
mcgs
spls
icgs-p
ifrms
sspls-d
sspls-v
prcus-a
prcus-i
prcus-p
prculs-v
prculs-d
pos
DP1
DP2
DP3
DP4
DP5
DP6
DP7
DP8
DP9
DP10
DP11
DP12
DP13
DP14
DP15
DP16
DP17
DP18
DP19
DP20
DP21
DP22
DP23
DP24
DP25
DP26
DP27
DP28
DP29
DP30
DP31
DP32
DP33
DP34
DP35
DP36
DP37
DP38
DP39

**Figure S2b. Manual PMC sulcal labels for every participant with developmental prosopagnosia (DP)**
Each sulcus is displayed on the left and right hemisphere inflated cortical surfaces in FreeSurfer 6.0.0, with each label displayed as an outline according to the key at the top. Each hemisphere contains at least 8 sulci (from anterior to posterior): ifrms, mcgs, spls, prcus-a, prcus-i, prcus-p, prculs-d, pos. An additional 4 sulci are variably present: icgs-p, sspls-d, sspls-v, and prculs-v.
